# Affinity-guided design of PDZ-binding peptides in a shared latent space

**DOI:** 10.64898/2026.11.11.765864

**Authors:** Iori Yamahata, Teppei Shimamura, Shuto Hayashi

## Abstract

PDZ-binding motifs (PBMs) are short C-terminal sequences that bind PDZ domains and contribute to protein localization and complex assembly. Designing PBMs could modulate these interactions while leaving most of the original protein sequence unchanged. Because PDZ recognition preferences overlap, modulating intended interactions requires considering off-target binding alongside target affinity. Computational PBM design methods generally use sequence-based affinity prediction and structure-based evaluation, but it remains challenging to guide sequence searches toward different binding objectives with a single trained model. Here, we propose a deep learning-based method to design PBMs by analytically formulating single-target, off-target-aware, and dual-target design objectives as directions in a PDZ–PBM shared latent space learned through quantitative PDZ–PBM affinity prediction. In single-target PBM redesign for DLG4-PDZ3, our method identified 100 top-ranked candidates, 84 of which were in the top 1% for experimentally measured interaction fitness in a published mutant library, compared with 64 using an existing method. In single-target *de novo* design of 10-residue PBMs, our method achieved the highest median *in silico* binding scores among the compared methods for three of the four PDZ domains tested. In off-target-aware and dual-target PBM design for PTPN3 and SCRIB-PDZ3, off-target-aware design increased between-group score differences within each domain at the cost of lower target-domain scores, whereas dual-target design yielded scores close to those of the respective single-target designs for both domains. Together, these results show that shared latent representations learned from quantitative affinities allow a single trained model to guide PBM design across different binding objectives without retraining.

## Introduction

Protein–protein interactions play important roles in regulating molecular localization and signal transduction within cells. Some of these interactions arise from the binding of short amino acid sequences to domains that recognize them. PDZ domains are one such class of recognition domains and primarily bind C-terminal sequences known as PDZ-binding motifs (PBMs). PDZ– PBM interactions support associations between receptors and scaffold proteins at postsynaptic sites [1] and protein assemblies at cell–cell junctions [2]. Recognition preferences overlap among PDZ domains, allowing the same PBM to bind different PDZ domains [3]. Designing PBM sequences could therefore adjust which PDZ domains a protein binds and the strength of those interactions. Such designs could provide tools for investigating and controlling localization and complex assembly while leaving most of the protein sequence unchanged. Sequence-based affinity prediction has previously been applied to PDZ–PBM design. Models predicting dissociation constants or binding free energies have been reported [4–6]. These predictions have supported experimental validation of designed high-affinity peptides [4] and searches for new ligand candidates [6]. Beyond improving affinity, design may require avoiding other PDZ domains with overlapping recognition preferences or promoting binding to multiple PDZ domains. Off-target-aware design has been demonstrated using sequence models that approximate structure-derived scores to optimize affinity and specificity [7]. Applying this approach to an additional PDZ domain requires structure-based calculations and construction of an approximation model for that domain. A method that searches for PBM sequences under target and off-target specifications without building a dedicated model for each domain would facilitate design across diverse PDZ domains.

In this study, we focus on a shared latent space in which affinity is represented by the relationship between target and candidate vectors. Similarity in shared latent spaces has been used for candidate selection [8] and domain– peptide binding prediction [9]. BALM, for protein–small-molecule interactions, and BALM-PPI, for protein pairs, independently encode each partner and learn to map cosine similarity to measured affinity [10, 11]. Here, we use this mapping both to evaluate candidates and to define sequence-search objectives. When predicted affinity increases linearly with cosine similarity under a common prediction equation, optimal directions in continuous space can be derived analytically. These directions maximize single-target predicted affinity, the predicted affinity difference between a target and one off-target, or mean predicted affinity for two targets. Using these directions as search objectives allows the same procedure to identify PBM candidates for different target and off-target specifications without retraining.

We propose a method that uses shared latent representations learned from quantitative PDZ–PBM affinities to search for PBM sequences under different design objectives. We define objective directions in latent space and search for sequences whose representations approach them. We first assess affinity prediction using sequence-similarity-aware cross-validation and external measurements. For single-target design, we redesign the four C-terminal positions of a known PBM and retrospectively compare the candidates with a published mutant library to confirm that the search yields candidates with high measured interaction fitness. We then apply unrestricted 10-residue design to multiple PDZ domains and compare the candidates with those from existing methods using structure prediction and interaction-energy calculations. We also perform off-target-aware and dual-target designs by switching objective directions within the same trained model. Comparisons with single-target designs show that objective-specific candidate selection is also reflected in calculated interaction properties.

## Methods

The method combines PDZ–PBM affinity prediction with sequence search using the learned representations. Two sequences are mapped into a shared latent space, where their similarity is trained to correspond to measured affinity. The trained model is then fixed, and PBM sequences are sought whose representations approach a direction determined by the target and off-target specifications.

### Training data and preprocessing

We use PDZ–PBM interaction data obtained by Gogl and colleagues using double-affinity purification and fluorescence readout, termed DAPF [3]. The dataset contains 10-residue PBMs composed of the 20 standard amino acids. After excluding eight missing observations, 57,581 observations across 133 PDZ domains are retained. Dissociation constants *K*_*d*_ are converted to p*K*_*d*_ = *−*log_10_(*K*_*d*_*/*M), so that higher values indicate stronger binding.

Measured p*K*_*d*_ values are used for the 17,437 observations with quantified affinities. The remaining 40,144 observations represent interactions too weak to quantify within the assay range. For these observations, a PBM-specific upper bound *L* on p*K*_*d*_ is calculated from the detection threshold and conversion equation in the original study. They enter training as censored observations satisfying p*K*_*d*_ *≤ L*, rather than as fixed affinity values.

### Shared latent representations for affinity prediction

A dual encoder represents PDZ and PBM sequences independently in a shared latent space (Fig. 1a). The model comprises a PDZ encoder, a PBM encoder, and a prediction head that maps their representations to affinity.

**Figure 1.**
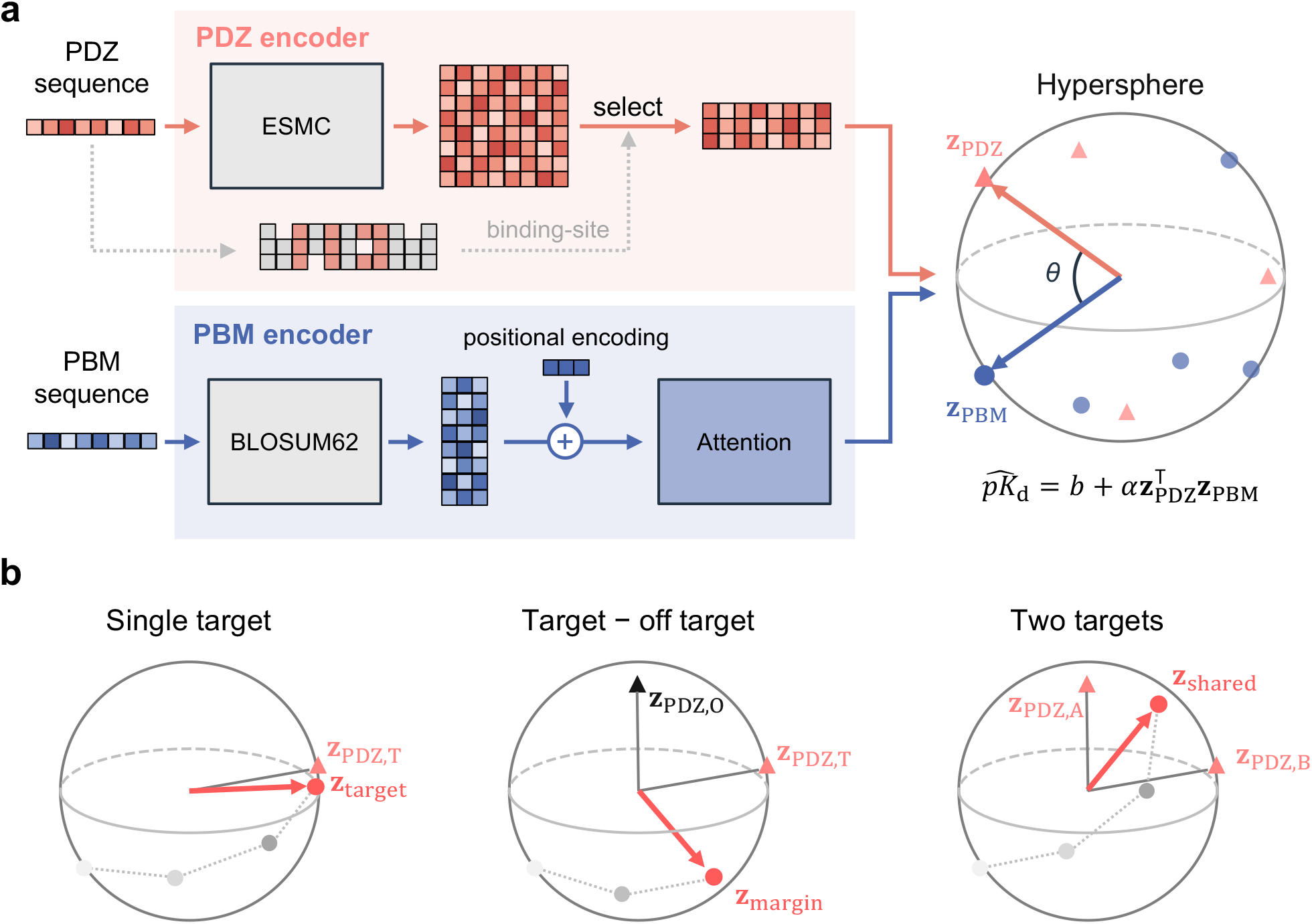
Overview of PDZ–PBM affinity prediction and sequence design. a, Model architecture. Full PDZ sequences are passed through ESMC-600M, and representations at 17 binding-site positions mapped through a reference alignment are extracted and concatenated. PBMs are encoded using position-specific BLOSUM62 features and a Transformer. Both outputs are projected to 128 dimensions and L2-normalized, and their inner product predicts p*K*_*d*_. b, Objective-specific directions. Single-target design follows the target vector *z*_target_. Design maximizing the predicted target–off-target affinity difference follows the difference vector *z*_margin_. Design maximizing mean predicted affinity for two targets follows the sum vector *z*_shared_. The spheres schematically represent the shared 128-dimensional unit hypersphere.

The PDZ encoder receives full domain sequences through the protein language model ESMC-600M [12], producing a sequence-length-by-1,152 feature matrix. Representations are extracted at 17 positions corresponding to the PBM-binding site. To identify these positions, input sequences are added with MAFFT [13] to a published multiple sequence alignment of 209 PDZ domains [14]. Residues are mapped to the 17 positions defined by Tonikian and colleagues from contacts in nine complex structures [15]. The position-specific representations are concatenated and mapped from 19,584 to 128 dimensions through LayerNorm, dropout, and a linear layer.

The PBM encoder represents each of the 10 C-terminal residues as a 20-dimensional vector based on BLO-SUM62 [16]. Each position is projected to 16 dimensions, combined with a learnable positional embedding, and passed through a Transformer layer. The representations of all 10 positions are concatenated and linearly projected into the same 128-dimensional latent space as the PDZ representations.

The prediction head L2-normalizes both encoder outputs. Let *z*_PDZ_ and *z*_PBM_ denote the resulting unit vectors, and let *s*_PDZ_ and *s*_PBM_ denote the input sequences. Predicted affinity is defined as

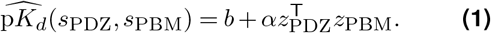

Here, *∥z*_PDZ_*∥* = *∥z*_PBM_*∥* = 1. The trainable parameters *a* and *b* are shared across all PDZ–PBM pairs. The positive coefficient *α* = softplus(*a*) scales similarity to affinity, and *b* is the intercept. Because the inner product of unit vectors equals cosine similarity, more closely aligned PDZ and PBM vectors yield higher predicted affinity.

### Loss function and model training

A Huber-based loss jointly fits quantitative affinities and weak-binding censoring constraints. Let *Q* and *C* denote the sets of quantitative and censored observations, respectively. For observation *i*, let *ŷ*_*i*_ be the prediction, *y*_*i*_ the quantified measurement, and *L*_*i*_ the censoring bound. The loss is

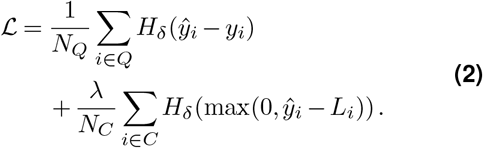

Here, *N*_*Q*_ and *N*_*C*_ are the numbers of quantitative and censored observations. The Huber loss *H*_*δ*_ increases quadratically for small errors and linearly for larger errors. Quantitative observations contribute their prediction errors, whereas censored observations contribute only the amount by which the prediction exceeds *L*_*i*_. Each term is averaged over its respective observation set.

The quadratic-to-linear transition is set to *δ* = 1, and the censored-loss weight is *λ* = 2.

Model parameters are optimized using AdamW.

### Single-target objectives and sequence search

For a target PDZ domain *T*, let *s*_PDZ,*T*_ and *z*_PDZ,*T*_ denote its sequence and latent vector. To increase predicted affinity for this domain, Eq. (1) serves as the sequence-search objective. Because *α* is positive, maximizing predicted affinity is equivalent to maximizing 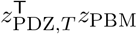. If the PBM unit vector can vary freely in continuous space, this inner product is maximized when it aligns with *z*_PDZ,*T*_ . The single-target direction is therefore defined as *z*_target_ = *z*_PDZ,*T*_ .

The model encodes sequences into latent representations but has no decoder that directly converts a target point into a sequence. We therefore fix the trained encoders, encode actual PBM sequences, and evaluate their similarity to *z*_target_. Sequence optimization uses beam search, which generates single-residue substitutions from the sequences retained at each step. Candidates are ranked by the objective, and a fixed number of top candidates are retained for the next step. Repeating this procedure searches for PBM sequences whose representations approach the target direction.

### Objectives with off-target or dual-target specifications

To increase the predicted affinity difference between a target *T* and one off-target *O*, the search objective is

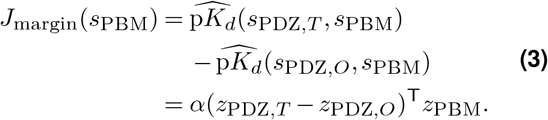

When *z*_PDZ,*T*_ ≠ *z*_PDZ,*O*_, the direction maximizing this objective on the unit hypersphere is

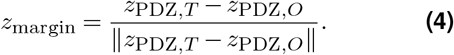

The maximum in continuous space is *α ∥z*_PDZ,*T*_ *−z*_PDZ,*O*_ *∥*. Replacing the target direction with this difference direction prioritizes the predicted affinity difference from the off-target.

When two PDZ domains, *A* and *B*, are equally weighted as targets, the objective is their mean predicted affinity:

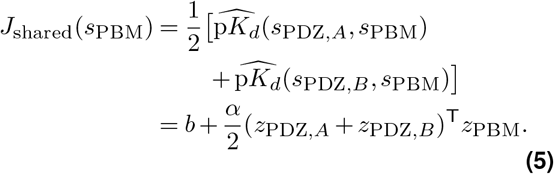

When *z*_PDZ,*A*_ ≠ *−z*_PDZ,*B*_, the maximizing direction on the unit hypersphere is

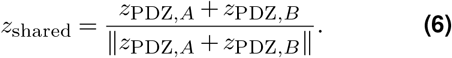

This objective increases mean predicted affinity without imposing equality between the two target affinities.

The target, difference, and sum vectors thus define the directions for single-target, single-off-target, and dual-target-mean objectives, respectively (Fig. 1b). The same beam-search procedure is used to search sequences for each objective.

## Results

### Affinity prediction from shared latent representations

We trained shared PDZ and PBM representations on the DAPF affinity data. Their arrangement is shown in Fig. 2a. We evaluated generalization to unseen sequences, the contribution of the PBM input window, and affinity ranking in an external dataset.

**Figure 2.**
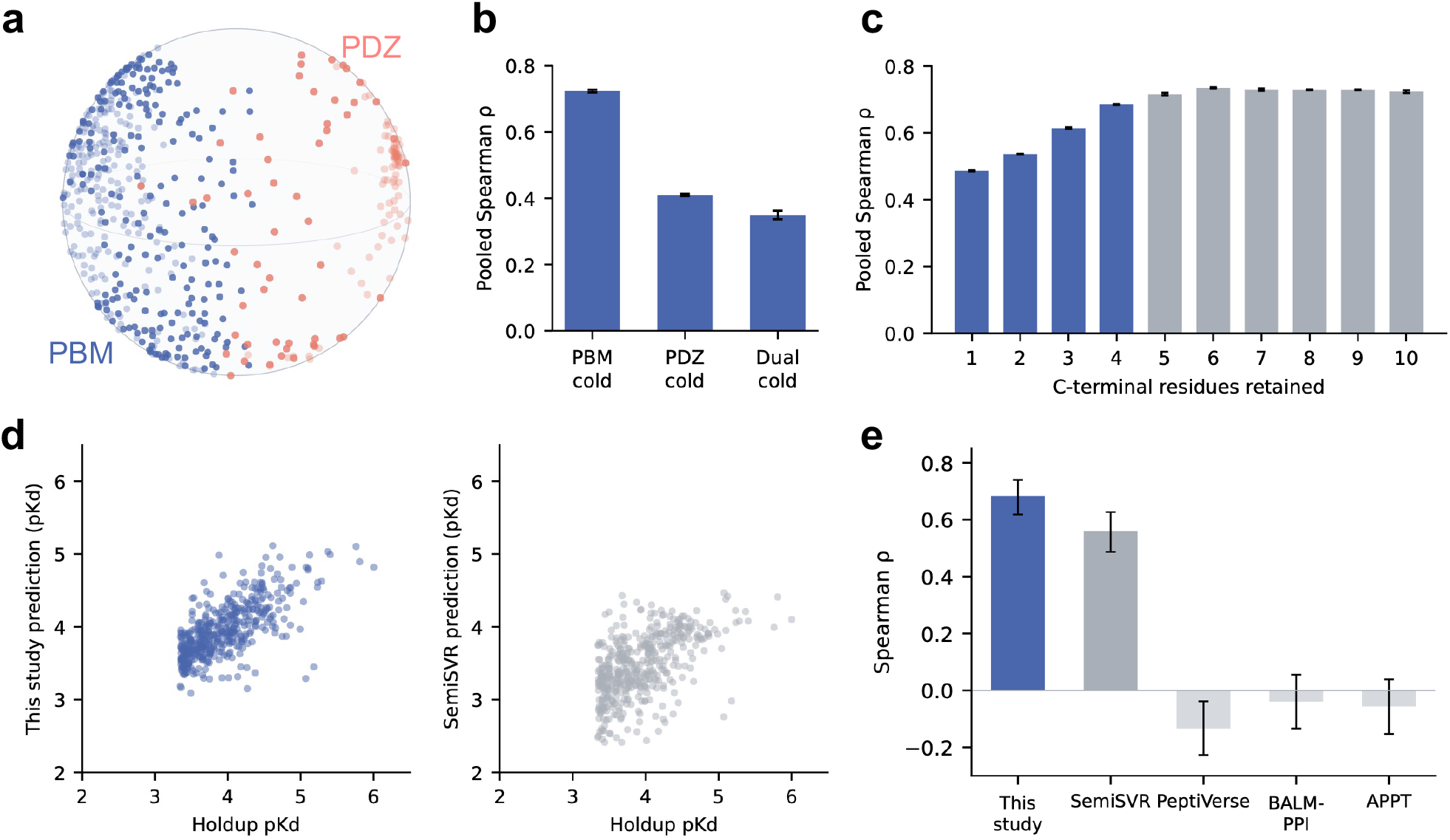
Shared latent representations and affinity prediction. a, Spherical multidimensional scaling (MDS) of 133 PDZ domains and 433 PBM records from one model trained with a PBM group held out. Angular distances between the original latent vectors are approximated on a three-dimensional unit sphere. Coral and blue circles indicate PDZ domains and PBMs, respectively. b, Pooled Spearman correlations across all 17,437 quantitative pairs under PBM-cold, PDZ-cold, and Dual-cold evaluation. c, The same metric under PBM-cold evaluation with one to 10 C-terminal input positions. One to four positions are shown in blue and five to 10 in gray. Bars and error bars in b and c show means and sample standard deviations across three training runs. d, Measured versus predicted affinities for 448 external DLG1-PDZ2 peptides, using our model (left) and SemiSVR (right). e, Spearman correlations for five predictors on the same 448 sequences. Our model uses the mean prediction of three trained models. Error bars in e indicate 95% intervals from 2,000 sequence-level bootstrap samples. Our method is labeled “This study”.

Under PBM-cold evaluation, which tests generalization to unseen PBMs, the pooled Spearman correlation across all quantitative pairs was 0.724 *±*0.004. The correlation was 0.410*±* 0.003 under PDZ-cold evaluation, where PDZ domains were excluded from training. It decreased to 0.350*±* 0.013 under Dual-cold evaluation, where both PDZ and PBM groups were excluded (Fig. 2b). Correlations were therefore lower when the PDZ domain itself was unseen than when new PBMs were evaluated against known PDZ domains. To assess PBM ranking for individual PDZ domains, we also calculated within-PDZ correlations. Their mean under PBM-cold evaluation was 0.461 *±*0.021.

Residues upstream of the four C-terminal positions can contribute to PDZ recognition [15, 17]. To test whether this information improves affinity prediction, we retrained models with different numbers of C-terminal input residues. Spearman correlation increased from 0.685 *±*0.001 with four positions to 0.735*±* 0.002 with six positions. With all 10 positions, the correlation was 0.724*±* 0.004 (Fig. 2c). Adding upstream information improved prediction over the four-position input, whereas extending the input from six to 10 positions provided no further improvement.

We evaluated affinity ranking for external peptides without training-set overlap using Holdup Multiplex measurements for DLG1-PDZ2 [18]. Figure 2d compares predictions from our model and the PDZ–peptide affinity regression model SemiSVR [5] for 448 sequences shared across five predictors. Spearman correlations were 0.683 for our model and 0.561 for SemiSVR. PeptiVerse, which targets protein–peptide interactions [19], and the protein-pair models BALM-PPI [11] and APPT [20] did not yield positive rank correlations on this sequence set (Fig. 2e).

### Single-target sequence search and comparison with experimental measurements

We assessed the effectiveness of single-target search by redesigning CRIPT for DLG4-PDZ3. Models were trained without the PBM group corresponding to the parent sequence. The six upstream residues were fixed, and the four C-terminal positions were searched. Candidates were retrospectively matched to published Binding-PCA measurements for 101,553 sequences [21]. BindingPCA fitness measures interaction-dependent yeast growth rather than dissociation constants. We therefore evaluated candidate ranks within the measured set rather than agreement in absolute values with predicted p*K*_*d*_.

Beam search from the parent sequence STKNYKQTSV selected STKNYKETAV as the highest-ranked prediction. This sequence also had the highest measured fitness among the 101,553 sequences and thus ranked first experimentally (Fig. 3a).

**Figure 3.**
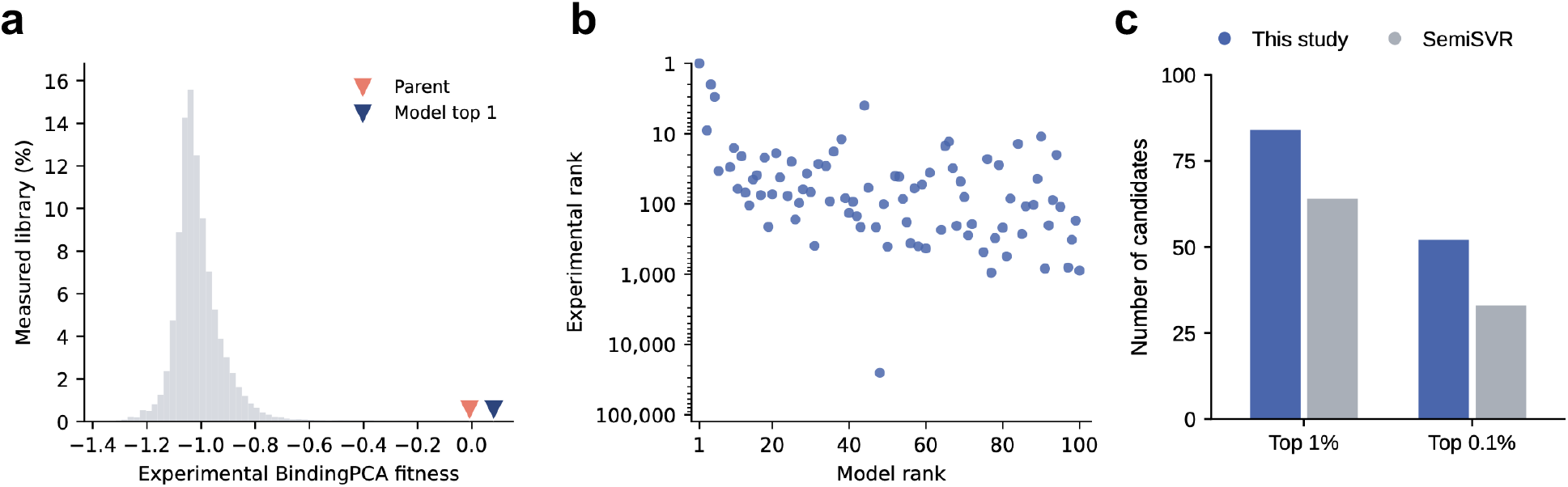
Single-target sequence search and external interaction measurements. a, BindingPCA fitness distribution for 101,553 published sequences. The parent `STKNYKQTSV` and top predicted candidate `STKNYKETAV` are marked near the horizontal axis by pink and navy downward triangles, respectively. b, Rank correspondence for all 85 measured candidates among our top 100 predictions. The horizontal axis shows predicted rank among the 1,024 searched sequences, and the logarithmic vertical axis shows experimental rank among 101,553 measured sequences. The 15 unmeasured candidates are not plotted. Experimental ranks are based on point estimates of fitness. c, Numbers of candidates confirmed to rank within the experimental top 1% or 0.1% among each method’s top 100 predictions. The top 1% and 0.1% thresholds are defined using the measured set, including the parent sequence.

To determine whether high-fitness recovery extended beyond the top candidate, we examined the top 100 predictions among the 1,024 sequences evaluated during search. Measurements were available for 85 candidates, whereas 15 were unmeasured. Of the 85 measured candidates, 84 ranked within the experimental top 1%, and 52 ranked within approximately the top 0.1%. However, the candidate ranked 48th by the model ranked 25,330th experimentally. Thus, the top predictions were concentrated in the high-fitness region, although substantial rank discrepancies remained for individual candidates (Fig. 3b). We compared our search with SemiSVR-guided search from the same parent sequence under the same budget of 1,024 evaluated sequences. Among the top 100 predictions, 84 candidates from our method and 64 from SemiSVR were confirmed to fall within the experimental top 1%. The corresponding counts for the top 0.1% were 52 and 33 (Fig. 3c).

### *In silico* evaluation of new 10-residue candidates

We extended single-target search to all 10 positions without fixing a terminal motif. The targets were DLG4-PDZ3, DLG1-PDZ2, SYNJ2BP, and SNX27. For each target, we evaluated 100 candidates from our method, SemiSVR-guided beam search, the language model PepMLM [22], and the diffusion model PepCCD [23]. Evaluation combined complex prediction using AlphaFold 3 [24], energy minimization with the ff14SB force field [25], and MM/PBSA interaction-energy calculations [26]. Details are provided in the Supplementary Information. *In silico* binding scores were obtained for all 1,600 candidates across the four targets and four methods.

For all four targets, mean cosine similarity between the retained sequences and the target direction increased during search and approached a plateau (Fig. 4a–d, left). Our candidates had the highest median scores among the four methods for DLG4-PDZ3 (Fig. 4a), DLG1-PDZ2 (Fig. 4b), and SYNJ2BP (Fig. 4c). Median differences from SemiSVR-guided search were 1.48, 1.08, and 4.11 kcal/mol, respectively. PepMLM had higher medians than PepCCD among the two generative models. Differences between our method and PepMLM were 8.64, 2.34, and 5.02 kcal/mol, respectively, with the magnitude varying across targets. For SNX27, the PepMLM median of 39.14 kcal/mol exceeded the median of 38.32 kcal/mol for our method (Fig. 4d).

**Figure 4.**
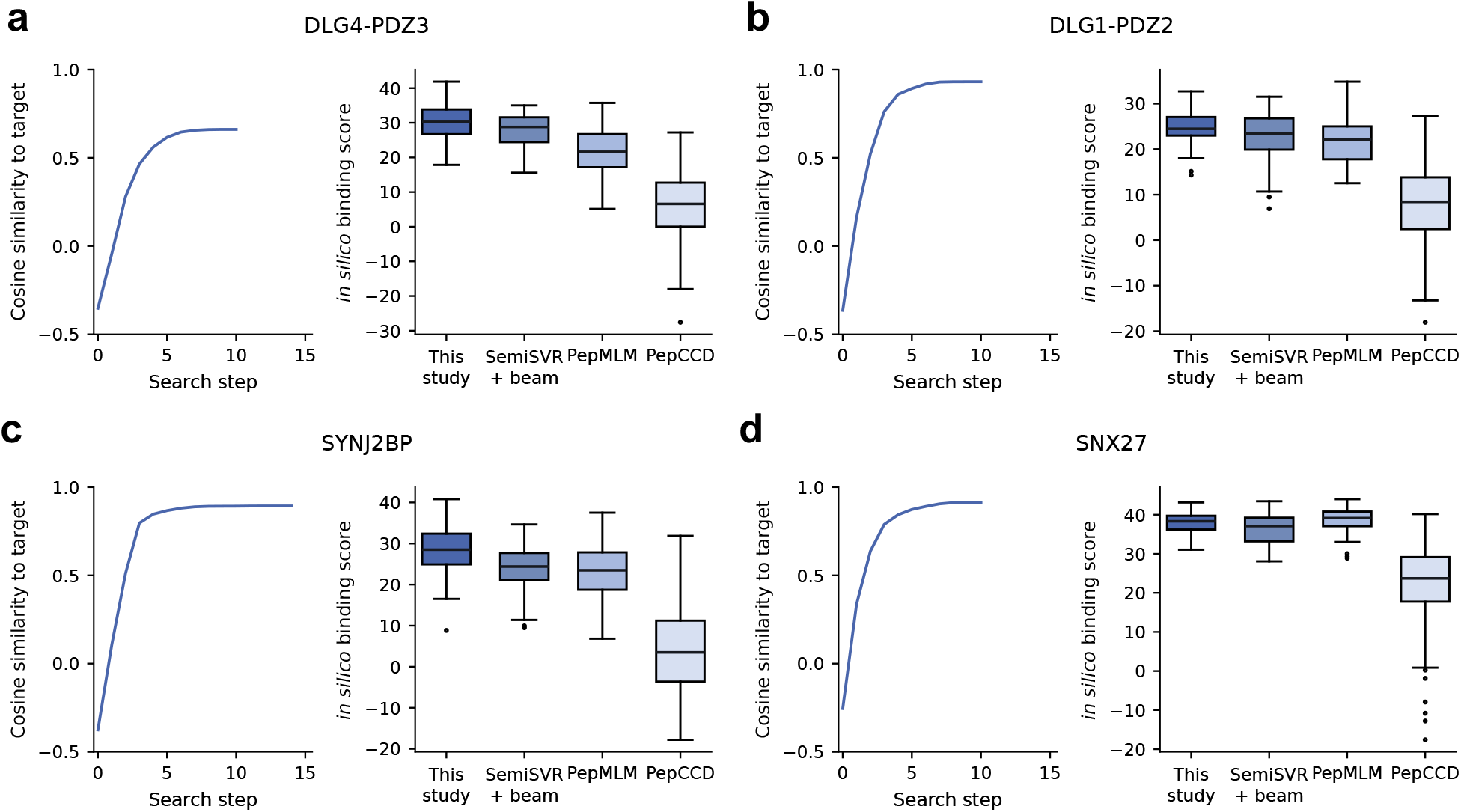
Single-target sequence search and structure-based evaluation of new candidates. In each panel, the left plot shows mean cosine similarity to the target direction across the 100 sequences retained at each search step. The right plot shows *in silico* binding scores for 100 candidates proposed by each method. a, DLG4-PDZ3. b, DLG1-PDZ2. c, SYNJ2BP. d, SNX27. “This study” denotes our method, and “SemiSVR + beam” denotes beam search using SemiSVR affinity predictions as the objective. Each group contains 100 unique sequences fixed before structural calculations. One predicted structure per candidate was energy-minimized to obtain the single-frame MM/PBSA score *S* = *−*ΔTOTAL. Higher scores indicate more favorable calculated interactions. Boxes show the first to third quartiles, central lines show medians, whiskers extend to values within 1.5 interquartile ranges, and points indicate outliers. Comparisons are made within the same PDZ domain.

None of the 400 sequences proposed by our method exactly matched a 10-residue sequence in the DAPF training data. However, the median nearest-neighbor Hamming distance within each target-specific candidate group was one residue, indicating that candidates were concentrated among closely related sequences.

### Changes in candidate groups under different target specifications

We next examined how candidate groups changed when prioritizing target affinity, the target–off-target affinity difference, or mean affinity for two targets. PTPN3 and SCRIB-PDZ3 have both been reported to recognize class I terminal motifs [27, 28]. We compared five conditions: each domain as the sole target, each domain as the target with the other as an off-target, and both as targets.

With the trained model and search settings held constant, we switched the objective direction among the target, difference, and sum vectors. Under all five conditions, mean cosine similarity between the 100 retained sequence representations and the objective direction increased during search and approached a plateau (Fig. 5a).

**Figure 5.**
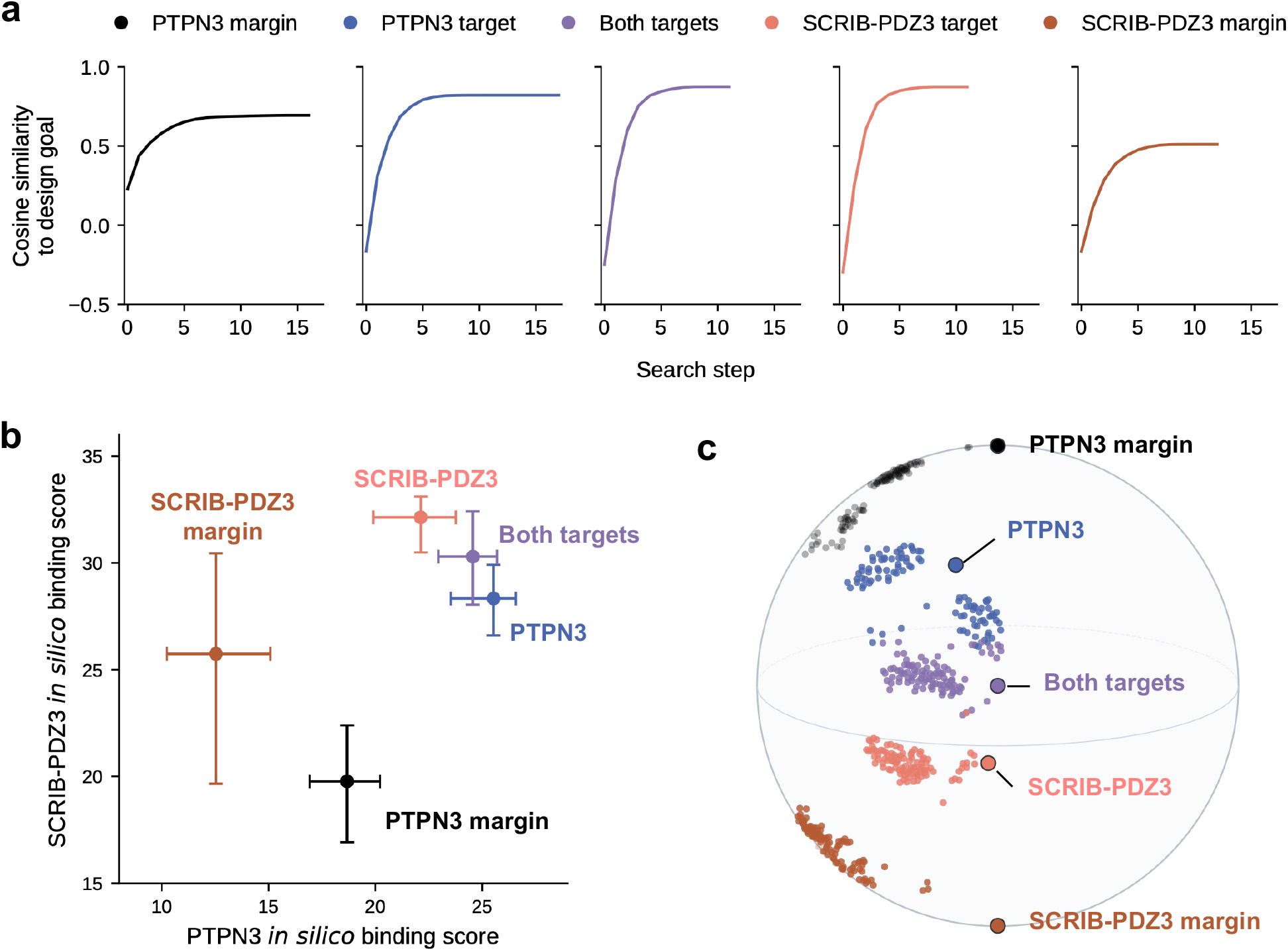
Candidate groups obtained by switching design objectives. “Target” denotes single-target design, whereas “margin” specifies the other PDZ as an off-target. “Both targets” maximizes mean predicted affinity for both PDZs. Colors are consistent across panels. a, Mean cosine similarity to the objective direction for the 100 sequences retained at each step, averaged across three models and five searches. b, Median *in silico* binding scores and interquartile ranges for 100 candidates per condition. Each sequence–PDZ score is the median from three structures; higher scores indicate more favorable calculated interactions. Comparisons are made within each PDZ. c, Spherical MDS of candidates and objective directions based on shared latent representations. Small and large circles indicate candidates and objective directions, respectively.

To assess whether design objectives also affected structure-based evaluations, we calculated *in silico* binding scores against both PDZ domains for the 100 candidates from each condition. Candidate groups were compared separately for each PDZ domain (Fig. 5b). Comparing the two single-target groups, the PTPN3-directed group had a 3.24 kcal/mol higher median score against PTPN3. Against SCRIB-PDZ3, the SCRIB-PDZ3-directed group had a 3.77 kcal/mol higher median score. For both domains, the group designed for that domain scored higher than the group designed for the other domain.

When the other domain was specified as an off-target, the differences between group medians increased to 6.14 kcal/mol for PTPN3 and 5.97 kcal/mol for SCRIB-PDZ3. However, the target-domain median decreased from 25.37 to 18.68 kcal/mol for the PTPN3-directed group and from 32.14 to 25.74 kcal/mol for the SCRIB-PDZ3-directed group. Off-target specification therefore reduced target-domain scores while increasing the difference between design groups within each PDZ domain. For dual-target design, median scores were 24.57 kcal/mol for PTPN3 and 30.30 kcal/mol for SCRIB-PDZ3. For each domain, the median lay between those of the two single-target groups. Unlike designs prioritizing the off-target difference, dual-target designs yielded scores close to those of each domain’s own single-target group.

The 100 candidates from each condition occupied different regions when their shared latent representations were visualized by spherical multidimensional scaling (MDS; Fig. 5c).

## Discussion

We used shared latent representations learned from quantitative PDZ–PBM affinities to search PBM sequences under different design objectives. The method represents single-target predicted affinity, the target–off-target predicted affinity difference, and mean predicted affinity for two targets as directions derived from one prediction equation. Changing the design objective requires neither model retraining nor candidate-specific complex calculations during search. Target and off-target specifications can therefore be changed within the same candidate-proposal procedure.

The learned representations supported both affinity ranking and the proposal of candidates with high measured interaction fitness. For external measurements of DLG1-PDZ2, the rank correlation exceeded that of SemiSVR (Fig. 2d,e). In CRIPT redesign, the highest-ranked candidate had the highest measured fitness among 101,553 sequences. Of the top 100 candidates, 85 had measured values, and 84 of these ranked within the top 1% experimentally (Fig. 3). The highest-ranked CRIPT candidate obtained by this search retained the same class I motif as the parent. Both sequences shared Thr at P*−* 2 and Val at the terminus. PDZ sequence preferences depend on positions beyond P*−* 2 and the terminal residue used in conventional class definitions [15]. These results suggest that the search distinguished sequences that cannot be separated by motif class alone and selected candidates with high measured fitness. In addition, unrestricted 10-residue design yielded higher median *in silico* binding scores than the comparison methods for three of the four targets (Fig. 4). Candidate proposals beyond partial redesign of a known PBM were therefore also supported by computational evaluation.

We also examined the effects of switching design objectives using PTPN3 and SCRIB-PDZ3. Off-target-aware design increased calculated score differences between candidate groups within each PDZ domain compared with single-target design. In contrast, dual-target designs yielded scores close to those of each domain’s own single-target group for both PDZ domains (Fig. 5). Differences between design objectives were reflected not only in affinity predictions but also in structure-based evaluations. Selecting objective directions without changing the model can therefore produce candidate groups for different design requirements.

Explicit objective directions are useful not only for candidate selection but also for monitoring search progress. For unit vectors that can vary freely in continuous space, cosine similarity to the objective direction reaches its maximum of 1 when the directions coincide. This upper bound provides a reference for assessing how closely the retained candidates approach the objective direction, in addition to whether the objective value has stopped increasing. For DLG1-PDZ2, mean similarity among the retained candidates reached approximately 0.93 (Fig. 4b), whereas off-target-aware designs plateaued at lower similarities than single-target designs (Fig. 5a). However, these curves alone cannot distinguish whether a plateau below 1 reflects limitations of the search procedure or the attainable representations of actual sequences.

A limitation of this study is that the binding properties of newly designed candidates have not been validated experimentally. While structure prediction with AlphaFold 3 and single-frame MM/PBSA calculations provide a practical computational evaluation, they remain approximations of actual binding free energies. Accordingly, to determine whether score differences calculated using AlphaFold 3 and MM/PBSA are reflected in actual binding, affinities for both the target and off-target need to be quantified under the same measurement conditions and compared with the intended binding changes. Moreover, censored weak-binding observations provide upper bounds on p*K*_*d*_, leaving limited quantitative information for learning affinity differences among weak binders. Quantifying candidate affinities for both the target and off-target is therefore important not only for design validation but also for supplementing training information in this regime.

A possible extension is to broaden the design scope from PBMs to PDZ domains. Because the same prediction equation evaluates both PDZ and PBM representations, the approach could be extended to PDZ sequence searches for a fixed PBM or to co-design of both partners. PDZ design would also need to account for folding and preservation of existing functions in addition to binding. However, prediction performance was lower in the PDZ-cold and Dual-cold evaluations (Fig. 2b), so this extension requires validation and improvement of predictions for PDZ sequence changes. Introducing PBMs with experimentally characterized binding properties into proteins and examining changes in localization or interaction partners could help investigate the roles of individual PDZ interactions. Such modifications might also promote molecular recruitment to specific scaffold proteins or the formation of desired complexes. PBM introduction or replacement can adjust interaction partners while leaving most of the protein sequence unchanged and, following cellular validation, can be used in such designs.

Together, these results show that shared latent representations learned from quantitative affinities can support the proposal of PBM candidates that prioritize discrimination against an off-target or binding to two targets, in addition to stronger target binding.

## Data availability

The DAPF, Holdup Multiplex, and BindingPCA datasets used in this study are available from the original publications cited in this article. Code for affinity prediction and PBM sequence design is available at https://github.com/IoriYamahata/PBM_design.

## Acknowledgements

This study was carried out using the TSUBAME super-computer at Institute of Science Tokyo.

## Author contribution

I.Y. conceptualized the study, conducted the investigation, developed the software, prepared the visualizations, and wrote the original draft. All three authors contributed to the methodology and acquired funding. T.S. and S.H. supervised the study and reviewed and edited the manuscript.

