## Supplementary Information for "Affinity-guided design of PDZ-binding peptides in a shared latent space"

### PDZ–PBM affinity prediction and sequence design using latent representations

#### Cross-validation of affinity prediction

PDZ domains and PBM measurement records are each partitioned into five groups using GraphPart 1.0.2 [1]. For PDZ domains, similarity is calculated as the number of matching residues in pairs detected by MMseqs2 [2], divided by the longer sequence length. PBM partitioning uses the fraction of matching positions across all pairs of 10-residue sequences.

PBM-cold and PDZ-cold evaluations use fivefold cross-validation, holding out one PBM or PDZ group, respectively. Dual-cold evaluation tests all 25 combinations of the five PDZ groups and five PBM groups. Any observation containing either the test PDZ group or the test PBM group is excluded from training.

For each split, the non-test data are divided into training and validation subsets. The model is fitted on the training subset, and the epoch count with the lowest validation loss is selected. The training and validation subsets are then combined, and the model is trained again from scratch for the selected number of epochs. Performance is evaluated on the held-out test data, and the procedure is repeated three times using the same split.

AdamW uses a learning rate of 0.001, weight decay of $10^{-4}$, and a maximum gradient norm of 5. During epoch selection, training continues for at most 1,000 epochs. It stops if validation loss fails to improve by at least $10^{-4}$ for 100 epochs.

#### Ablation of input representations

The PDZ input is replaced with either a 1,152-dimensional mean of all residue-level ESMC representations or a 340-dimensional concatenation of one-hot encodings at 17 positions. Missing aligned residues are represented by zeros in the one-hot input.

For PBMs, the BLOSUM62 features and their LayerNorm and linear projection are replaced with a learnable 16-dimensional amino acid embedding. Positional embeddings and self-attention are retained. This modification is tested both alone and together with the 17-position PDZ one-hot input.

Replacing the 17-position ESMC input with the whole-domain mean or 17-position one-hot input yielded PBM-cold correlations of 0.713 and 0.709, respectively. These values were close to the full-model correlation of 0.724. In PDZ-cold evaluation, however, correlations decreased from 0.410 to 0.208 and 0.202. In Dual-cold evaluation, they decreased from 0.350 to 0.178 and 0.058 (Fig. S1). Changing the PDZ representation therefore had a greater effect when evaluation included unseen PDZ domains.

Replacing PBM BLOSUM62 features with learnable embeddings yielded correlations of 0.721 under PBM-cold and 0.408 under PDZ-cold evaluation, close to the full-model values. The Dual-cold correlation was 0.305, below that of the full model. Combining PDZ one-hot inputs with learnable PBM embeddings yielded correlations of 0.703, 0.199, and 0.075 for PBM-cold, PDZ-cold, and Dual-cold evaluations, respectively.

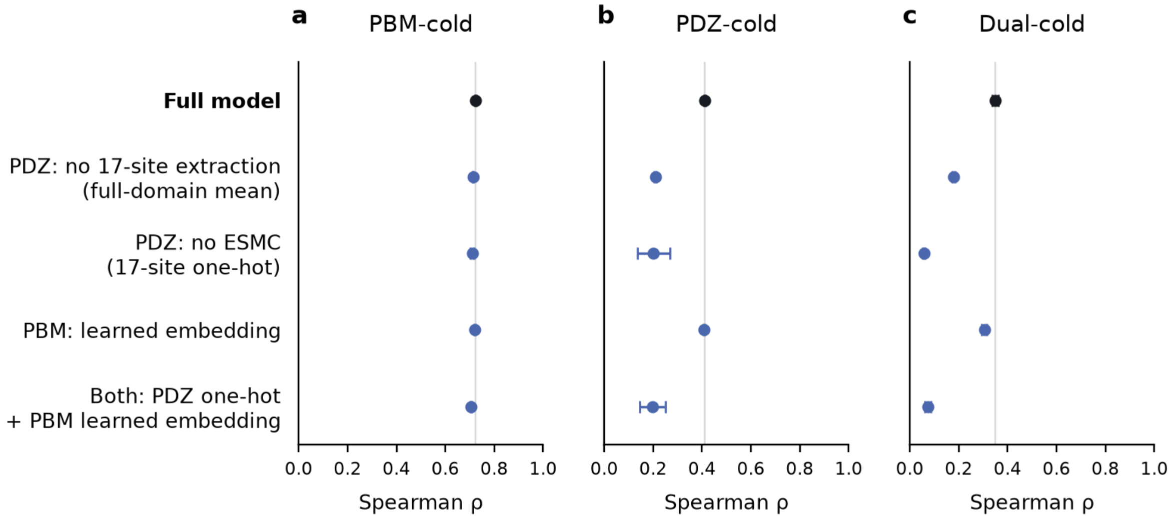


**Figure S1.** Ablation of input representations. a, PBM-cold. b, PDZ-cold. c, Dual-cold. Points show mean Spearman correlations across three training runs, and error bars indicate sample standard deviations. Black denotes the full model, blue denotes modified inputs, and gray vertical lines indicate the full-model means.

### Comparison of PBM input windows

For Fig. 2c, BLOSUM features outside the retained window are set to zero, preserving the 10-position input shape. Data splits are defined using the original 10-residue sequences. Models using one to nine C-terminal positions are each trained three times with the same splits and training settings described above for cross-validation. The 10-position condition uses models already trained under the same settings.

### Affinity ranking against external measurements

Sequences exactly matching DAPF sequences or the published SemiSVR training sequences are removed from the Holdup Multiplex data [3].

Holdup prediction and new 10-residue designs use three models with the same architecture, trained separately on all DAPF observations. Training duration is determined by inner validation under PBM-cold evaluation. External predictions are the mean predicted $\mathrm{p}K_d$ across the three models. SemiSVR [4] uses a Python port with the fixed parameters of the public MATLAB model. Publicly released PeptiVerse [5], BALM-PPI [6], and APPT [7] models are used without retraining.

### Sequence-search evaluation using a published mutant library

The N-terminal residues `ST` are added to the eight-residue sequences in the CRIPT library [8]. We exclude the 86 PBM measurement records in fold 4, which contains the DAPF sequence `DTKNYKQTSV` corresponding to the parent. Predictions use three models trained under the same conditions after this exclusion. The DLG4-PDZ3 sequence is taken from DAPF.

Our method and SemiSVR use the same parent sequence and variable positions, with a beam width of 4, at most 64 steps, and a budget of 1,024 unique evaluated sequences, including the parent. Candidates are selected in order of predicted binding strength; our method ranks them by decreasing predicted $\mathrm{p}K_d$. SemiSVR candidates are ranked using prediction scores based on the logarithm of the dissociation constant $K_d$.

### Design of new 10-residue sequences

Beam search for our method and SemiSVR uses 100 randomly generated starting sequences, a beam width of 100, and at most 250,000 unique evaluated sequences per search. Search terminates when the top 100 retained sequences no longer change or when the next candidate expansion would exceed the evaluation budget. For our method, the objective is computed separately by each of the three models. Candidates are ranked by the mean of these scalar objective values.

In the method comparison, our method and SemiSVR use the same starting sequences. Candidates are selected by decreasing predicted $\mathrm{p}K_d$ for our method and increasing predicted log-$K_d$ score for SemiSVR. PepMLM-650M generates 1,000 sequences with top-$k = 3$ and temperature 1.0, and candidates are selected by increasing conditional pseudo-perplexity. PepCCD uses the public weights and official sampler for 1,000 generation runs, each comprising 1,000 steps, with candidates retained in generation order. Each method contributes 100 unique 10-residue sequences composed of the 20 standard amino acids.

For the comparison of design objectives, five searches are performed with different starting sequences. Within each search, the starting sequences are shared across all conditions. The top 100 candidates from each search are pooled and deduplicated, and the 100 candidates with the highest mean objective are selected.

### Complex prediction and interaction-energy calculations

AlphaFold 3 [9] uses the officially distributed model weights. The same saved multiple sequence alignment is used for each PDZ domain, while no alignment is supplied for the peptide. Structure templates are not used, and the number of recycles is 10.

Energy minimization is performed with GROMACS [10], using heavy-atom positional restraints of $1{,}000$ kJ mol$^{-1}$ nm$^{-2}$. Minimization uses at most 5,000 steps and a force-convergence threshold of $100$ kJ mol$^{-1}$ nm$^{-1}$. Both peptide termini are free and ionized; cysteine is reduced, and histidine protonation is assigned according to the structure.

The gmx_MMPBSA [11] settings are an internal dielectric constant of 2, an external dielectric constant of 80, and a salt concentration of 0.15 M. Temperature is 298.15 K, with `PBRadii=4`, `radiopt=0`, and `inp=2`.

### Visualization of shared latent representations

Figure 5c uses one model trained on all DAPF observations. Spherical MDS minimizes the mean squared difference between all pairwise angular distances in the original 128-dimensional representation and the three-dimensional unit-sphere configuration. The configuration is displayed using orthographic projection.
